# Deletion of endothelial KLF4 as a model for preeclampsia

**DOI:** 10.64898/2026.03.30.715448

**Authors:** Emily Meredith, Andrew T. Meredith, Arya Mani, Martin A. Schwartz

**Affiliations:** Yale Cardiovascular Research Center, Section of Cardiovascular Medicine, Department of Internal Medicine, School of Medicine, Yale University, New Haven, CT 06511, USA; Department of Genetics, Yale University, New Haven, CT 06510, USA; Department of Cell Biology, Yale University, New Haven, CT 06510, USA; Department of Biomedical Engineering, Yale University, New Haven, CT 06510, USA

**Keywords:** Hypertension, gestational hypertension, sFLT1, preeclampsia, endothelial cell, KLF2/4

## Abstract

Preeclampsia (PE), or gestational hypertension, affects around 5% of pregnancies and leads to approximately 70,000 maternal and 500,000 fetal deaths per year worldwide, with increased cardiovascular and metabolic disease in survivors. PE is associated with elevated circulating levels of the alternative splice isoform of VEGF receptor 1 (sFlt1), defects in placental vasculature, kidney damage and, in severe disease, fetal growth restriction. Current mouse models induce PE via direct expression of sFlt1 or elevation of blood pressure, which bypass the natural risk factors for human disease, such as age, obesity, hypertension and diabetes. These risk factors have in common reduced expression of Krüppel-like factors 2 and 4 (KLF2/4), the endothelial transcription factors that protect against cardiovascular disease. We now report that inducible deletion of *KLF4* in maternal endothelium (*KLF4*^iECKO^) results in gestational hypertension, elevated sFlt1, defective placental vasculature, kidney damage and fetal growth restriction. *KLF4*^iECKO^ may thus serve as a mouse PE model suitable for mechanistic analysis and screening of treatments that address upstream risk factors.

## Introduction

Despite advances in detection and classification, preeclampsia (PE) remains a major cause of fetal death and maternal morbidity and mortality worldwide [1, 2]. In addition to short term effects on health, women after preeclamptic pregnancy show large increases in cardiovascular disease (CVD) incidence and mortality [3]. CVD risk scales with PE severity [4], defined as late (> 34wks) vs early-onset (< 34wks), as mild (140mmHg < BP < 160mmHg) vs severe (BP > 160mmHg), and with vs without fetal growth restriction (FGR)[4]. The most dangerous form of PE is early-onset, severe, with FGR, which raises the incidence of maternal CVD later in life as well as the incidence of metabolic syndrome and neurodevelopmental delays in the child [5]. Methods to manage PE are limited, indeed, the only cure is delivery of the placenta and fetus. Management is possible, but treatment options are limited out of concern for the developing fetus.

The principal risk factors for PE – age, obesity, hypertension and diabetes – mirror those for CVD. Widely used PE mouse models induce hypertension by infusion of angiotensin II or elevation of sFLT1 through viral overexpression, bypassing the endogenous regulatory pathways that govern PE (ref?). Thus, they do not account for the upstream risk factors in human disease. These human risk factors, however, share a common feature: they are opposed by endothelial cell (EC) expression of Krüppel-like factors 2 and 4 (KLF2/4), homologous transcription factors with overlapping (but nonidentical) gene targets and functions in ECs [6-8]. Klf2/4 expression in ECs declines with age [9, 10] and in diabetes [11-13] including gestational diabetes [14]. Elevating Klf2 or 4 confers resistance to multiple CVDs in mouse models [15, 16]. KLF2/4 function is vital to blood pressure regulation by inducing eNOS to control vascular tone and limiting vascular inflammation[17, 18]. While mouse EC-specific knock out (ECKO) of all four alleles of Klf2 and 4 is lethal, ECKO of 1-2 copies of these genes did not affect mouse survival [17]. A recent study showed that ECKO of Klf4 had little effect in young mice but markedly accelerated vascular aging (Jain, soon, I hope).

Measurement of blood pressure and circulating sFLT1 is the current standard for diagnosing PE, while sFlt1 is also critical to its pathophysiology [19, 20]. The placenta is thought to be the main source of circulating sFLT1, though a contribution from the maternal vascular has not been ruled out and could explain features of disease progression [21, 22]. Here, we report the development of a genetic model of PE driven by EC-specific KLF4 deletion, which offers a physiologically relevant platform that more closely mimics the human risk factors that drive PE [12].

## Methods

### siRNA KD of KLF2 or 4 in HUVECs

HUVECs were plated such that they were between 50-80% confluent at the time of siRNA treatment. Lipofectamine RNAiMAX (ThermoFisher Cat #13778075) was used to transfect 10nM KLF2, KLF4 or control siRNA into cells in Opti-Mem, as recommended by the manufacturer. The mixture was incubated on cells overnight in complete media (Lonza EGM-2 Endothelial Cell Growth Medium-2 BulletKit cat #CC-3162) before they were replenished and allowed to recover for 24h prior to collection for qPCR analysis.

For all qPCR analysis, cDNA was generated using the iSCRIPT cDNA Synthesis Kit (BioRad Cat#1708890) and qPCR was performed using SSO Advanced SYBR Green (BioRad Cat#1725270). Primer sequences used are as follows:

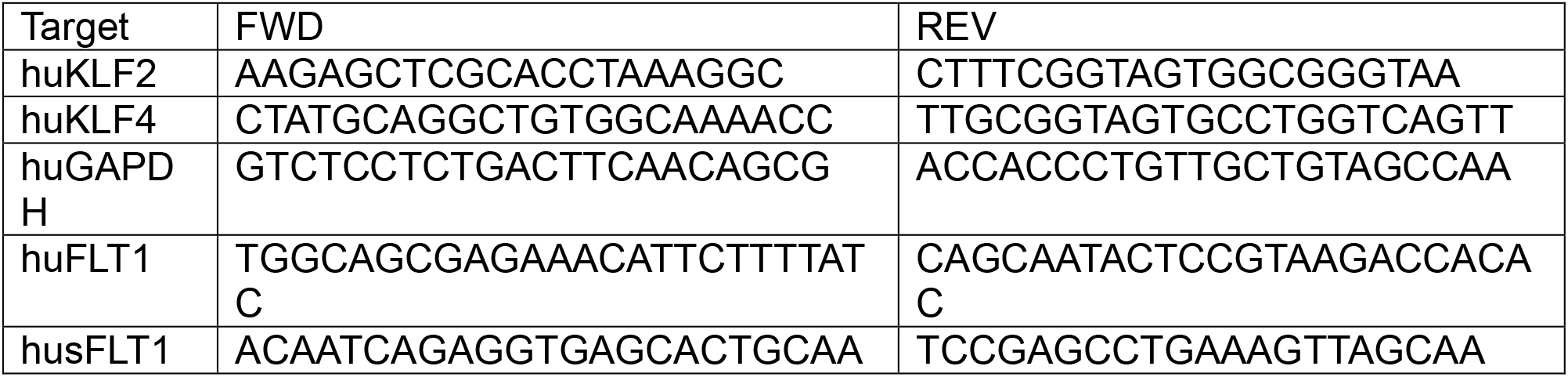

### RNAscope on Placenta Sections

Placentas were dissected at GD18.5, separated from the yolk sac and pup and immersion-fixed in 4% PFA overnight at 4C with gentle rocking. Images were taken of the underside (pup side) for visceral placenta vascularization analysis. Placentas were then incubated in 30% sucrose overnight at 4C with gentle rocking. Once placentas were saturated, as indicated by their sinking in the sucrose solution, they were cut in half longitudinally and frozen cut side down in OCT at -80C. Rapid freezing was accomplished by using acetone chilled by dry ice. A cryostat was then used to cut 10um longitudinal sections.

Slides containing placenta sections were then baked at 60C for 1hr, washed in PBS and post-fixed in 4% PFA for 1hr. Slides were then dehydrated in EtOH prior to beginning H2O2 treatment. ACD’s Multiplex Fluorescent V2 protease-free protocol was followed. sFLT1 probe: FLT1 probe: multiplex kit:. Following completion of the hybridization protocol, sections were then subjected to FN protein staining. Briefly, sections were washed in TBS/0.1%Tween 20 (TBST) three times, 5mins each. The sections were then blocked in TBS/5%BSA for 1hr followed by FN antibody incubation (Millipore cat#F3648, 1:500) for 2h at RT. Primary antibody was then washed off with TBST and secondary antibody (Alexa Fluor series from ThermoFisher; 1:1000) plus DAPI (ThermoFisher Cat#D1306, 1:10,000) was added in TBS/5% w/v BSA for 1hr at RT. Sections were then washed a final time before using TrueView Autofluorescence Kit (Vector Laboratories cat#SP-8400-15) diluted 1:10 to decrease background staining. Slides were then mounted using Vector Lab’s VactaShield vibrance antifade mounting medium (cat#H-1700-2). Imaging was done on a Zeiss Leica Confocal Microscope using the Leica Application Suit X software. A 40x oil objective was used for all images. Z-stacks of approximately 10um were taken with at least 5-6 images taken per sample. With RNAscope, it can be challenging to identify real versus background signal. Two ways we distinguished signal from noise were by firstly setting imaging parameters based off sections that were stained with positive and negative control probes provided by ACD. Secondly, by utilizing z-stacks, areas that displayed real signal were more apparent if RNAscope spots appeared in multiple z-sections. Maximal projections were used to enhance spots that were real versus not real. Image analysis was done using Fiji with maximal z-projections.

### sFLT1 ELISA Measurements

Blood samples were taken from dams at GD18.5 via cardiac puncture. Blood was collected into EDTA-coated K2 tubes (VWR cat#76343-512) and spun at 10,000g for 10mins at 4C to separate the serum, which was then isolated and flash frozen in LN2 before being stored at -80C. Samples were then thawed on ice prior to analysis. sFLT1 serum levels were analyzed using R&D’s VEGR1 ELISA kit (Cat #MVR100) with the DuoSet ELISA Ancillary Reagent Kit (Cat #DY008B). Serum samples were diluted from 1:5-1:20 to ensure samples landed within the standard curve. Raw pg/mL values are plotted.

### Mice

#### iECKO and Timed matings

8-9wk old KLF4^f/f;CDH5cre-ERT2+^ dams were injected with 80mg/kg/day tamoxifen for 5 days to induce ECKO. Dams were rested for 1wk following injections as tamoxifen can reduce fertility [23]. At the same time, male B6 littermates were separated 1wk prior to combination with females. Dams were then provided with dirty male bedding for 72hrs prior to introduction of males to induce estrus. 1-2 females were then introduced to male cages for 3 days before separating. Pregnancy was confirmed via plug checking, 7-day weight gain ≥2g or by blood pressure status.

#### Blood pressure monitoring via the CODA System

Mice were trained for 3-5 days prior to data collection. Training consisted of regular blood pressure measurements within the system. To reinforce training, foraging mix (VWR cat #76628-302) or other treats were given following any blood pressure collection. Blood pressure was taken every other day for 2 weeks following confirmation of pregnancy (GD7.5-GD17.5). Data per day consists of at least 3 measurements averaged. Data for non-pregnant control mice consists of at least 3 averaged measurements every other day for 2 weeks giving at least 18 averaged measurements per control mouse. To quantify acquisition stability, we calculated the percent coefficient of variation (%CV) for repeated tail-cuff cycles within each session for each mouse. Early acclimation sessions exhibited higher variability, consistent with handling-related stress, whereas CV progressively decreased with training. Sessions demonstrating %CV values within the expected physiological range (<10%) were considered stable.

### Kidney Analysis

#### Sections and IF staining

Kidneys were isolated from perfusion-fixed mice at GD18.5 and incubated in 4% PFA overnight at 4C with gentle rocking. Kidneys were then soaked in 30% sucrose overnight at 4C with gentle rocking or until the kidney was saturated as indicated by it sinking in the solution. Kidneys were then cut in half longitudinally, dabbed of excess sucrose and embedded in OCT with the cut-side down. Rapid freezing was accomplished using acetone chilled with dry ice. 10um sections were taken using a cryostat set at -20C. For staining, slides were thawed at room temperature (RT) and washed three times in PBS for 5mins to remove excess OCT. Samples were then blocked and permeabilized in PBS/0.1% TritonX/5% BSA for 1hr at RT. Primary antibodies were diluted in PBS/TritonX/BSA (FN (Millipore cat#F3648, 1:500), CD31 (Fisher Scientific cat#AF3628, 1:1000), CD34 (Abcam cat#AB81289)) and incubated overnight in a humidity chamber at 4C. Primary antibodies were then washed off with PBS/0.1% tween20 (PBST) three times for 5mins. Alexa Fluor Secondary antibodies were diluted in PBS/TritonX/BSA at 1:1000 and left for 1hr at RT. Slides were then washed again in PBST before the excess was blotted off. Slides were mounted using Prolong Gold antifade mounting medium (Invitrogen cat#P36980) and allowed to cure overnight prior to imaging. Imaging was done on a Zeiss Leica Confocal Microscope using the Leica Application Suit X software. A 20x air objective was used for all kidney images. Image analysis was done using Fiji.

#### H&E staining

Kidney sections were taken as described in the previous section. H&E staining of tissue sections was done by the Yale Research Histology Core using standard techniques

#### Urine collection and ACR

Urine was collected from GD17-18.5 dams prior to dissection using a clean cage without bedding, covered with saranwrap. The mouse was allowed to freely walk around until naturally voiding at which point the urine was quickly collected via clean syringe and stored at -80C prior to analysis. ACR was then measured using two kits: Albuwell M (Ethos biosciences cat#CHM03V055, and the creatinine companion kit (Ethos biosciences cat# CHM03V058) Samples were diluted 1:2-1:5.

## Results

### KLF2 association with hypertension

Given that both PE and cardiovascular disease share key risk factors, we interrogated the CVD Knowledge Portal (CVDkP) for variants in KLF2/4. Although associations between KLF2/4 SNPs and PE did not reach statistical significance, likely because of the limited statistical power of the study, variant rs3745318 in KLF2 is associated with blood pressure and coronary artery disease (Figure 1A and [24]). Furthermore, this variant is identified in the GTEx database as an eQTL for KLF2 in whole blood (p = 1.27e-6), supporting a potential regulatory role in vascular pathways linked to PE (figure 1B).

**Figure 1.**
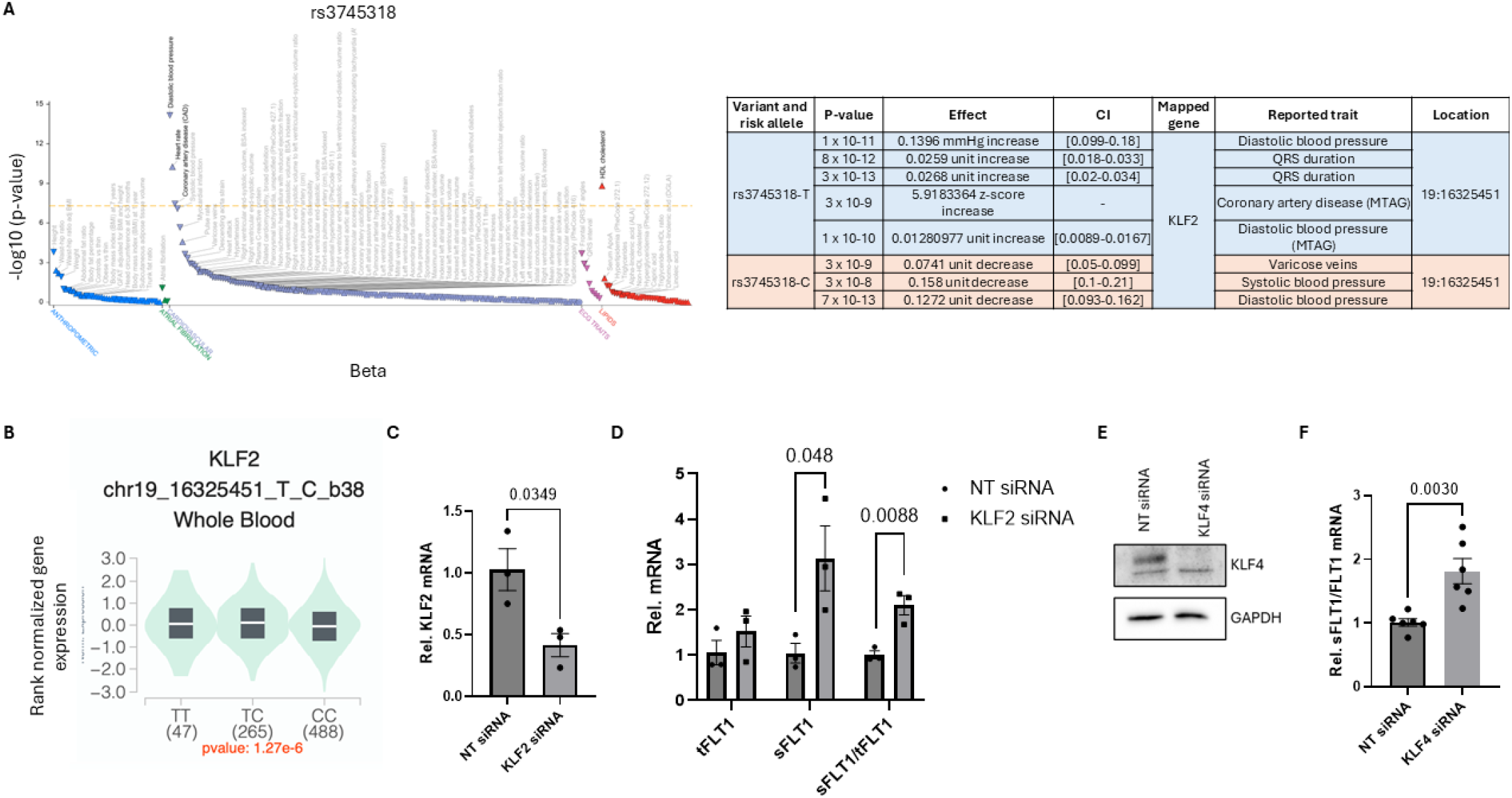
KLF2/4 are involved in hypertension and sFLT1 elevation. **A** CVD Knowledge Portal (CVDkP) analysis revealed SNPs within KLF2 linked to hypertension, a major causal factor in the development of PE. **B** This variant is identified in the GTEx database as an eQTL for KLF2 in whole blood. **C-F** siRNA-mediated KD of KLF2 (**B-C**) or 4 (**D-E**) in HUVECs significantly increased the sFLT1/FLT1 ratio in comparison to control siRNA. We did not observe a change in total FLT1 levels, indicating that KLF2/4 negatively regulates the splicing of sFLT1 and not the overall transcript levels of FLT1.

To address function, we suppressed KLF2 or 4 expression in HUVECs in vitro and used QPCR to quantify both total Flt1 (tFlt1) and the soluble isoform (sFlt1). siRNA-mediated knockdown (KD) of Klf2 (validated in Fig 1C) increased the sFlt1/tFlt1 ratio by approximately 2-fold, without significantly changing total Flt1 (tFlt1) mRNA (Fig 1D). Klf4 knockdown (validated in Fig 1E) similarly increased the sFlt1/tFlt1 ratio (Fig 1F). Knockdown of Klf2 or 4 thus induces a shift in mRNA splicing rather than expression. The established causal role of sFlt1 in PE, together with genetic links between Klf2/4 and hypertension, prompted us to examine a possible role for KLF signaling in PE [24].

### KLF4 ^iECKO^ triggers severe PE

In mice, early-onset, severe preeclampsia is characterized by elevated blood pressure (average BP>130) before the third trimester, elevated circulating sFLT1, and evidence of kidney or other organ damage [4, 25]. KLF4^f/f;CDH5Cre-ERT2^ dams were treated with tamoxifen to specifically delete endothelial KLF4 (KLF4^iECKO^) then bred with wild-type B6 males, producing WT offspring. Thus, any observed phenotype is attributable to the maternal genotype rather than the fetus.

Blood pressure was measured using a CODA tail-cuff telemetry system. To ensure accuracy, within-session variability, expressed as percent coefficient of variation (%CV) of tail-cuff measurements was calculated. %CV declined across acclimation sessions, marked on all graphs as a purple box, indicating effective mouse training (supplemental figure 1A-C). Percent CV was calculated per individual per experimental session and sessions above 10% CV were discarded (Supplemental figure 1A-C). We observed no difference in resting blood pressure between KLF4^iECKO^ and KLF4^f/f^ dams prior to the onset of pregnancy (Figure 2A, time point -1; supplemental figure 1D-F), as is often the case in human PE. However, GD6.5, KLF4^iECKO^ dams had a trend toward elevated BP by day 2.5, did not go through the mid-gestation nadir at around GD8.5, and remained elevated over controls at later times (Figure 2A; supplemental figure 1G-H). KLF4^iECKO^ thus elevates blood pressure within the first trimester of pregnancy in mice.

**Figure 2.**
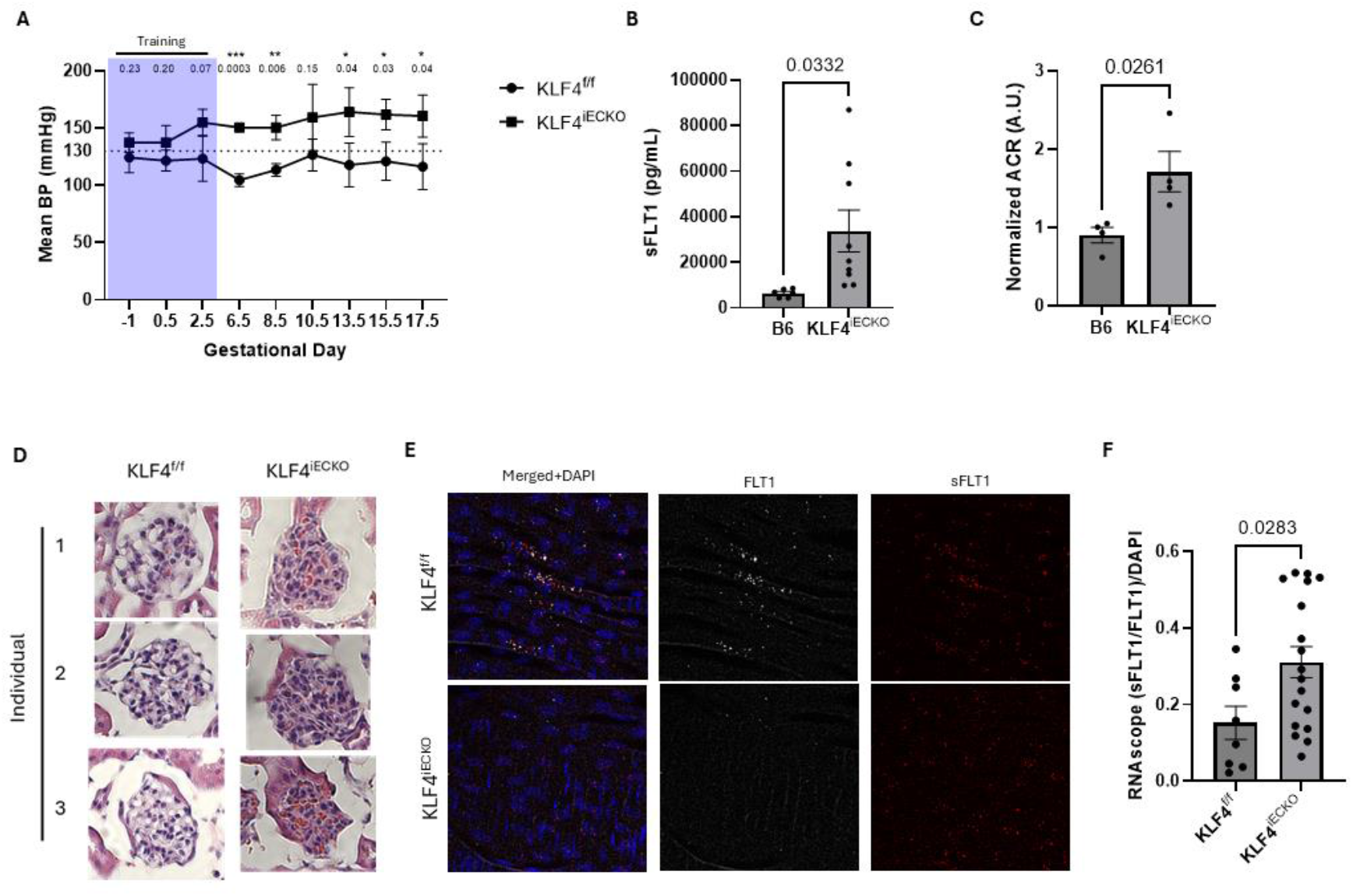
KLF4^iECKO^ mice develop early, severe preeclamptic pregnancies. **A** mean arterial pressure (MAP) was tracked using tail-cuff telemetry over the course of KLF4f/f or KLF4^iECKO^ pregnancies. Dams lacking EC KLF4 showed significant hypertension in comparison to cre-controls for the duration of their pregnancy. Prior to pregnancy, mice showed no difference between genotypes (GD-1). Days covered by purple are training days. **B** Blood samples were collected at GD18.5 via cardiac puncture. Samples were then run at dilutions ranging from 1:5-1:20 to ensure values landed within the standard curve. 6 B6 and 9 KLF4-/- dams were analyzed using R&D’s VEGFR1 ELISA kit. KLF4-/-mice had significantly higher sFLT1 in comparison to B6 controls. **C** To determine the extent of kidney damage during KLF4^iECKO^ preeclamptic pregnancies, we collected urine prior to sacrificing on GD18.5 and calculated the albumin:creatine (ACR). KLF4^iECKO^ dams had significantly more proteinurea in comparison to B6 controls. **D** This was further confirmed via kidney H&E staining, which showed severe occlusion of the glomeruli in KLF4^iECKO^ versus cre-controls. **E-F** RNAscope analysis of ECs in the aorta showed that KLF4^iECKO^ dams had significantly more sFLT1 splicing in their aortic endothelium in comparison to cre-animal controls.

To assess similarities to human preeclampsia, we next measured circulating sFLT1 and indicators of end organ damage. At GD18.5, pregnant KLF4^iECKO^ dams had strongly elevated sFLT1 compared to controls (Figure 2B; supplemental figure 2A). Kidney damage was analyzed using CD34 staining, a proteinuria assay and H&E glomeruli staining. These metrics revealed increased proteinuria, CD34 expression and capillary occlusion in KLF4^iECKO^ dams compared to controls (figure 2C-D, supplemental figure 1H-I) [26].

While evidence concerning which cell types contribute to elevated sFLT1 in the maternal circulation is mixed, it is known that maternal ECs can produce sFLT1 [27]. This source potentially creates a positive feedback loop, promoting maternal endothelial dysfunction and CVD risk, which can worsen fetal sFlt1 production. We therefore assayed sFlt1 mRNA in arterial endothelium of KLF4^iECKO^ dams using RNAscope probes specific for sFLT1 and total FLT1(supplemental figure 2B-D). Examination of aortic endothelium *en face* revealed elevated sFLT1 mRNA in ECs in KLF4^iECKO^ dams compared to cre-controls (Figure 2E-F). Together, these data indicate that KLF4^iECKO^ dams display a severe, early-onset PE phenotype.

### KLF4^iECKO^ triggers fetal growth restriction

Severe PE is associated with poor placental perfusion, leading to fetal growth restriction (FGR). Measures for FGR in mice include reduced pup weight and litter size, which were both strongly decreased in KLF4^iECKO^ litters (Figure 3A-B). Additionally, placental vascular area, measured both by longitudinal sections and macroscopically, was decreased in KLF4^iECKO^ placentas (Figure 3C-D). RNAscope analysis of placenta sections showed increased sFLT1 mRNA in the dense labyrinth in KLF4^iECKO^ placentas (Figure 3E; supplemental figure 3). Overall, these data confirm that maternal deletion of endothelial Klf4 results in severe preeclamptic pregnancy and FGR.

**Figure 3.**
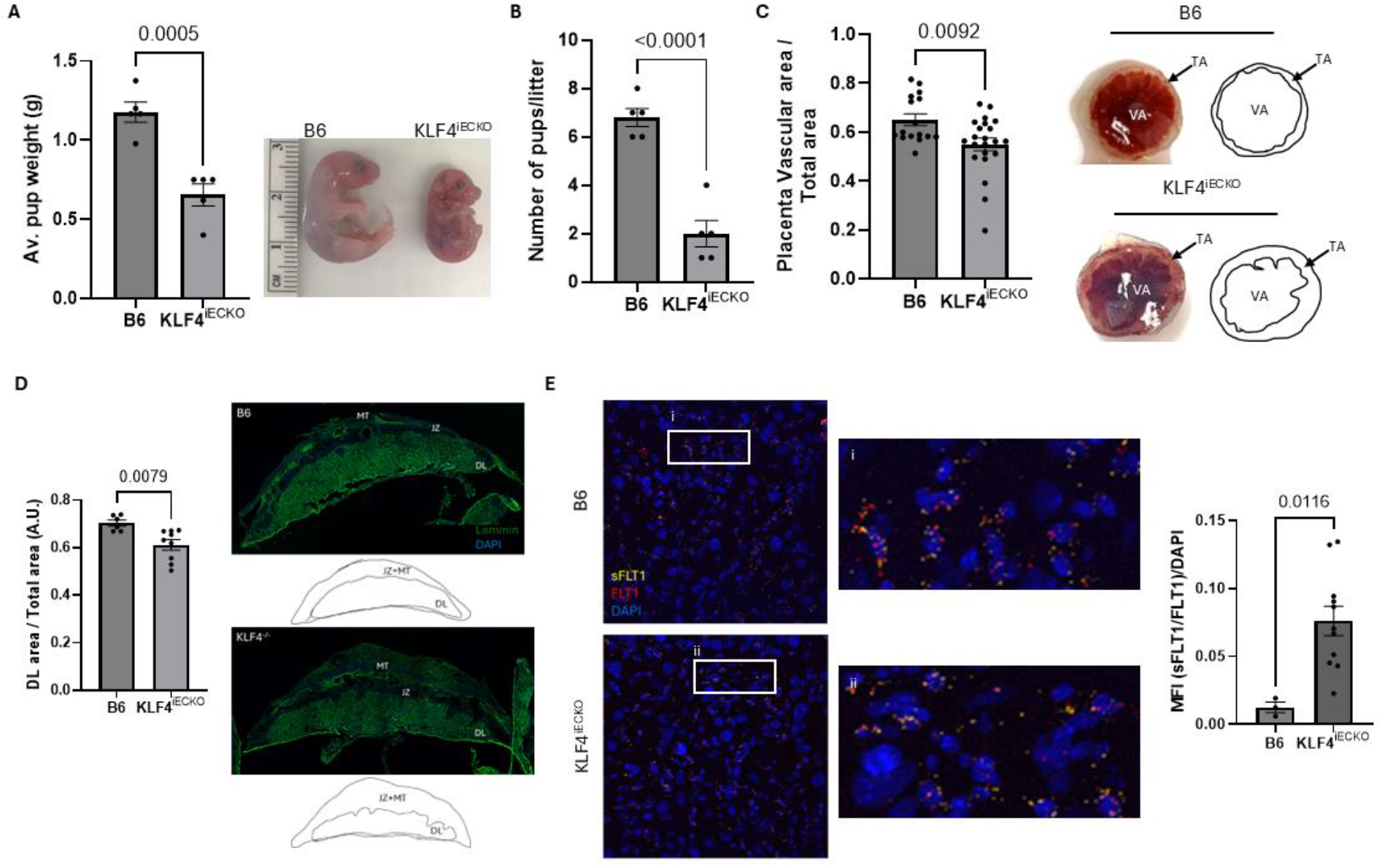
Pups from KLF4^iECKO^ Dams experience fetal growth restriction and decreased placental vascular area. **A** Following dissection from the yolk sac and placenta, pups were weighed as a group and the average pup weight was determined for each litter by dividing the total weight by the number of pups. Data points represent litters. Pups from KLF4^iECKO^ pregnancies were significantly smaller in comparison to B6 controls. A representative pup comparison is pictured to the right of the quantification. **B** In addition to being smaller, litters from KLF4^iECKO^ pregnancies had significantly fewer pups in comparison to B6 litters. **C-D** Placental vascular area was analyzed in two ways: firstly, via gross analysis of the underside (pup side) of the placenta where blood perfusion can be used to distinguish between the vascular area (VA) and the total area (TA) (**C**), and secondly via sectioning and comparing the dense labyrinth area to the total placental area (**D**). Representative images with area tracings are shown for both analyses. Gross images were taken using an iPhone 8. Placenta section images were taken using a 20x air objective with tiling used to capture the entire section. Data is represented as per placenta (**C**) with at least 6 sections being analyzed per genotype in 3 independent staining (**D**). **E** RNAscope analysis with FN protein staining on placenta sections reveals increased sFLT1 splicing and FN deposition in KLF4^iECKO^ versus B6 controls. Representative images are shown with zoomed insets (**Ei-ii**) and quantification. Data points are placentas that are an average of 3 stained sections, placentas from at least 3 different litters were used for each genotype. All data is shown as the mean +/- SEM.

## Discussion

Preeclampsia is a severe disease of pregnancy leading not only to maternal or fetal death but to life-long increased incidence of CVD in surviving mother [28, 29] and increased metabolic syndrome, impaired vascular function and neurodevelopmental problems in surviving offspring [5, 30]. Mild PE is associated with a ∼2-fold increased incidence of maternal CVD while severe, early onset PE increases future CVD by almost 10-fold [31]. Our data identify maternal KLF4^iECKO^ as a novel genetic model of early onset PE with FGR, the most severe form. Unlike viral sFLT1 overexpression, this approach preserves endogenous regulation and maternal–placental interactions, providing a physiologically relevant framework for studying how maternal vasculature defects contribute to development of PE. It may also better represent disease in humans where the main risk factors are known to decrease endothelial Klf2 and 4 expression. PE was also observed in a spontaneously hypertensive rat model [32] but the mouse KLF4^iECKO^ model offers greater opportunity for reverse genetic functional analysis as well as better recapitulating PE predisposition in otherwise healthy women [30].

Treating PE is a considerable challenge, due to a large extent to the risk of harm to the fetus. Hypertension management through aspirin administration is currently the only widely used therapy to mitigate PE symptoms. Our model suggests that the maternal vasculature may offer viable targets. Indeed, the KLF2/4 target gene eNOS, a major component of blood pressure regulation and target of hypertensive risk factors, is under consideration as a target for PE treatment [33, 34]. Recently developed antibodies [6] and siRNaS [35] that target vascular endothelium and elevate Klf2/4 expression, perhaps in conjunction with delivery via nanoparticles, Fab fragments and some antibody subtypes, e.g., IgA and IgM, that do not efficiently cross the placental barrier offer some promise here. The Klf4^iECKO^ model therefore offers a suitable platform to test these interventions in future studies.

## Acknowledgments

We thank the Yale Research Histology Core, M. Jain (Brown University) for the KLF4^f/f;CHD5cre-ERT2^ mice, H. Aldrich for sample OCT embedding, and the Schwartz Lab members for the extensive discussions.

## Figures

**Supplemental Figure 1.**
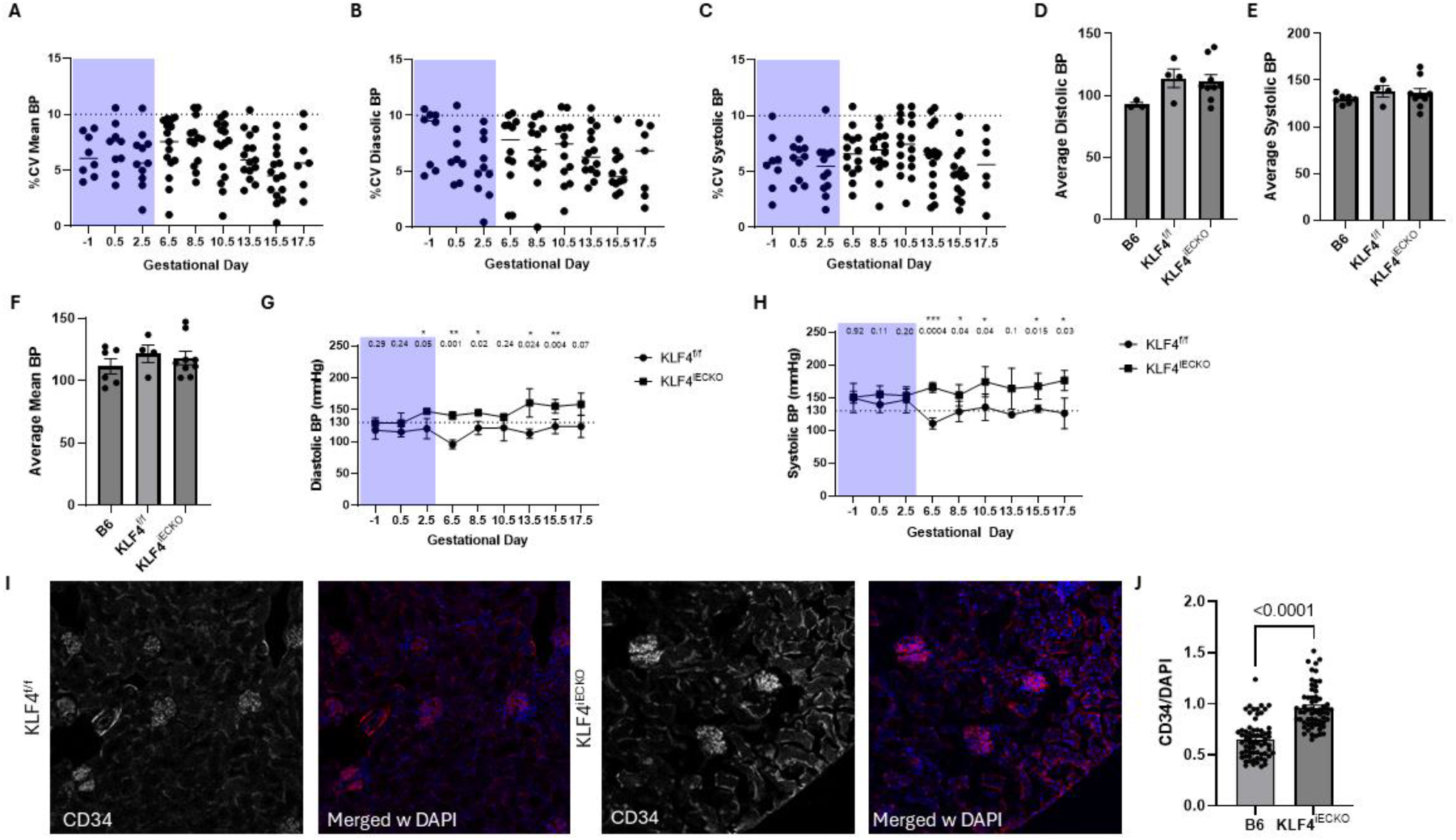
Additional assays on hypertension and kidney damage. **A-C** percent coefficient of variation (%CV) calculated per session per individual to ensure robust measurements. For each animal and session, BP was calculated as the mean of ≥3 accepted tail-cuff cycles. Session stability was quantified using the coefficient of variation (CV = SD/mean × 100). Cycle-level outliers exceeding ±2 SD were excluded to remove technical artifacts. Sessions demonstrating CV values within the expected physiological range (<10%) were considered stable. Days covered by purple are training days. **D-F** average diastolic (**D**) and systolic (**E**) and mean (**F**) blood pressure comparison between nonpregant B6, KLF4^f/f^ and iECKO dams showing no difference prior to pregnancy. **G-H** Time course of diastolic (**G**) and systolic (**H**) blood pressure changes during pregnancy in f/f or iECKO KLF4 animals. Days covered in purple are training days. **I-J** Representative CD34 staining in KLF4^iECKO^ and f/f controls with quantification (**I**).

**Supplemental Figure 2.**
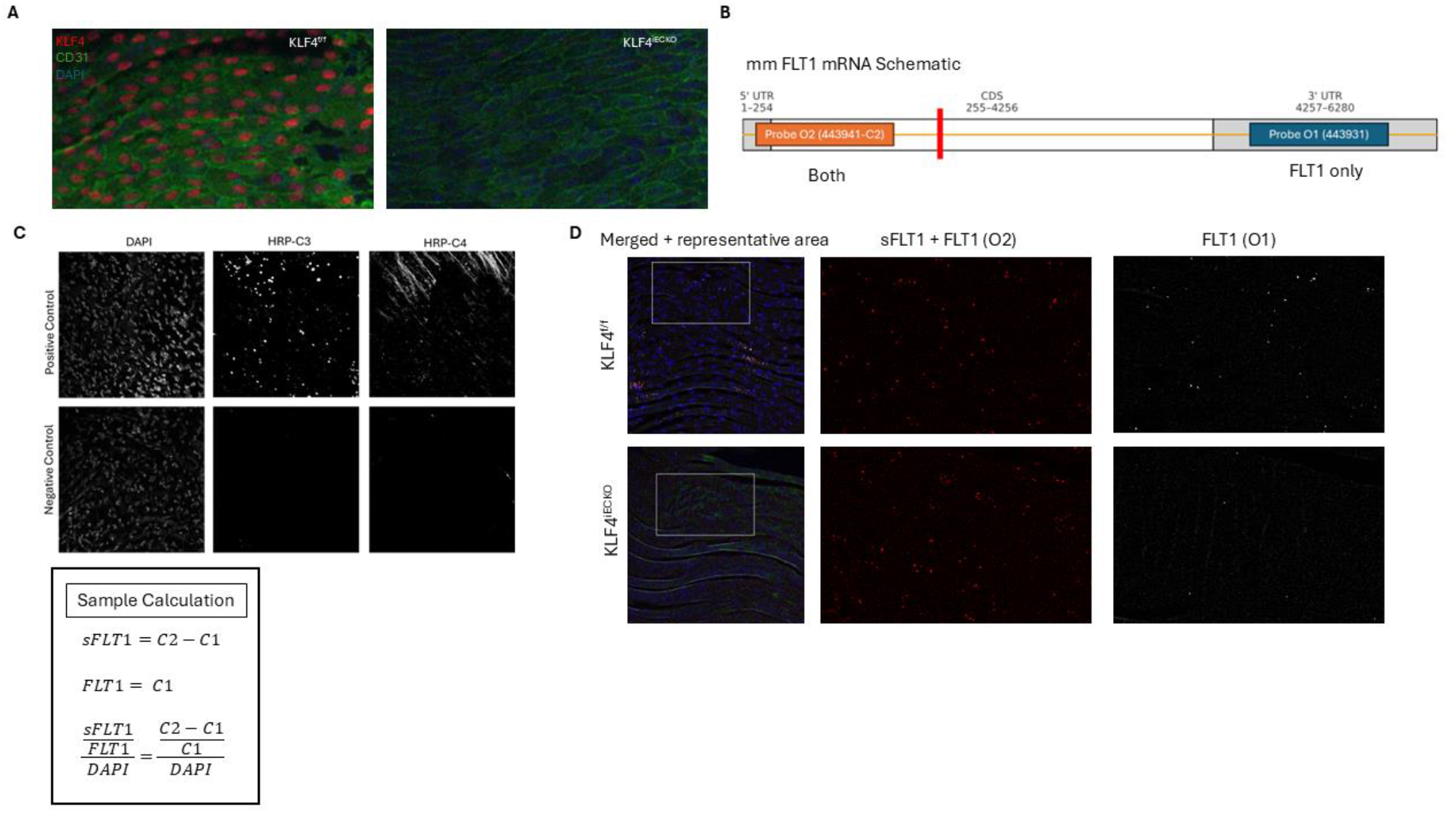
RNAscope Schematic and Analysis. **A** confirmation of KLF4^iECKO^ following tamoxifen injection. Aortas were stained *en face* for KLF4, CD31 and DAPI to confirm iECKO. **B** Mouse FLT1 mRNA schematic is shown with RNAscope probe locations and the sFLT1 i13 splice site (red line). **C** RNAscope probe control staining in *en face* aorta pieces. Laser power and exposure for each repeat experiment was determined by control probe fluorescence before imaging experimental samples. **D** Split channel representative RNAscope *en face* aortas with FN protein staining and sample calculation.

**Supplement Figure 3.**
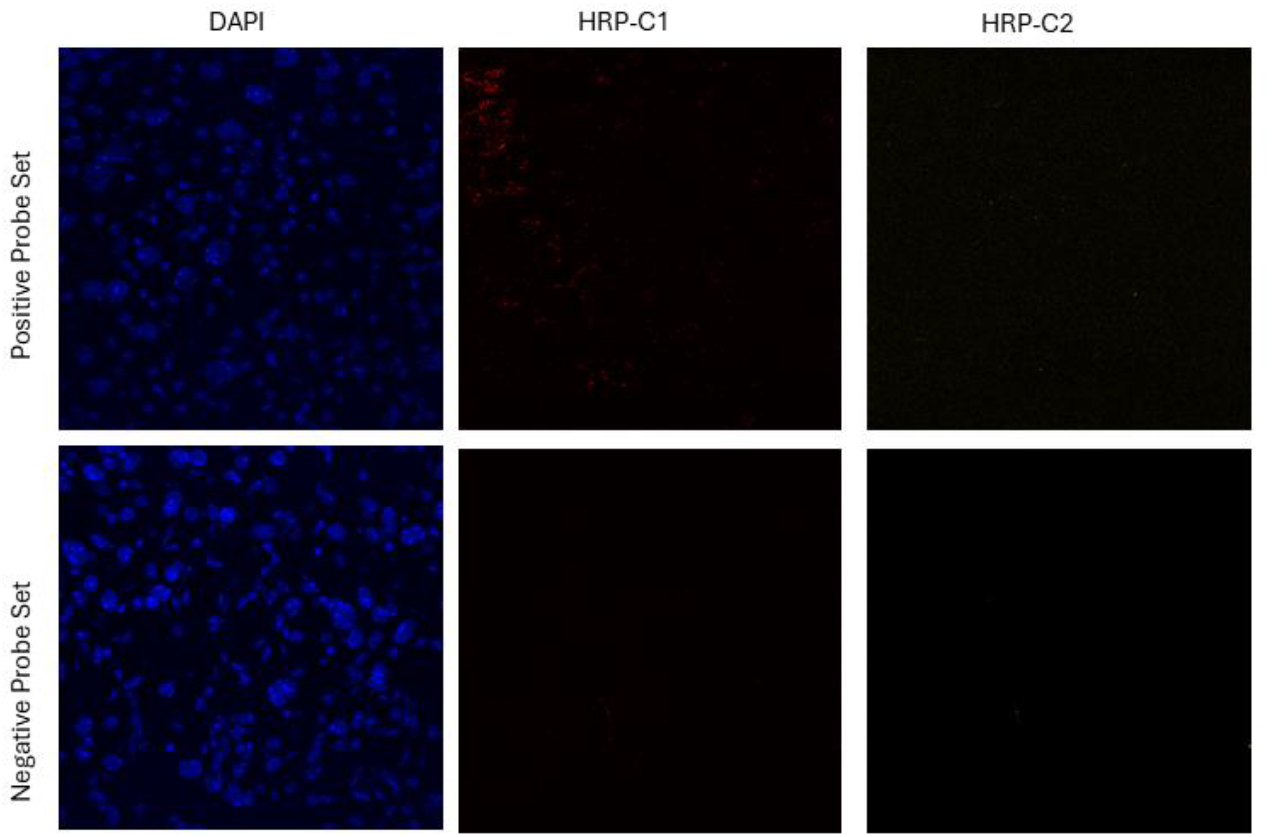
Placenta RNAscope validation using mouse positive and negative control probes. Placenta sections from 2-3 individual litters were used to stain for positive and negative control probes. Imaging parameters (laser power, exposure) for experimental samples were determined based on detection of signal only in the positive control set, but not the negative control set, as represented here. Channels HRP-C1 and –C2 were optimized in this experiment, we did not develop HRP-C3 since that channel was taken by FN protein staining.

## References

1. Cresswell, J.A., et al., Global and regional causes of maternal deaths 2009–20: a WHO systematic analysis. Lancet Glob Health, 2025. 13(4): p. e626–e634.

2. Ma’ayeh, M. and M.M. Costantine, Prevention of preeclampsia. Semin Fetal Neonatal Med, 2020. 25(5): p. 101123.

3. Bellamy, L., et al., Pre-eclampsia and risk of cardiovascular disease and cancer in later life: systematic review and meta-analysis. BMJ, 2007. 335(7627): p. 974.

4. Roberts, J.M., et al., Subtypes of Preeclampsia: Recognition and Determining Clinical Usefulness. Hypertension, 2021. 77(5): p. 1430–1441.

5. Huang, K.H., et al., Prediction of pre-eclampsia complicated by fetal growth restriction and its perinatal outcome based on an artificial neural network model. Front Physiol, 2022. 13: p. 992040.

6. Joshi, D., et al., Endothelial gamma-protocadherins inhibit KLF2 and KLF4 to promote atherosclerosis. Nat Cardiovasc Res, 2024. 3(9): p. 1035–1048.

7. Hsieh, P.N., et al., Aging and the Kruppel-like factors. Trends Cell Mol Biol, 2017. 12: p. 1–15.

8. Deng, H., et al., A KLF2-BMPER-Smad1/5 checkpoint regulates high fluid shear stress-mediated artery remodeling. Nat Cardiovasc Res, 2024. 3(7): p. 785–798.

9. Hsieh, P.N., et al., A conserved KLF-autophagy pathway modulates nematode lifespan and mammalian age-associated vascular dysfunction. Nat Commun, 2017. 8(1): p. 914.

10. Dykxhoorn, D.M., et al., MicroRNA-29c-3p and -126a Contribute to the Decreased Angiogenic Potential of Aging Endothelial Progenitor Cells. Int J Mol Sci, 2025. 26(9).

11. Samak, M., et al., Dysregulation of Kruppel-like Factor 2 and Myocyte Enhancer Factor 2D Drive Cardiac Microvascular Inflammation and Dysfunction in Diabetes. Int J Mol Sci, 2023. 24(3).

12. Thakar, S., et al., Intermittent High Glucose Elevates Nuclear Localization of EZH2 to Cause H3K27me3-Dependent Repression of KLF2 Leading to Endothelial Inflammation. Cells, 2021. 10(10).

13. Zhong, F., et al., Reduced Kruppel-like factor 2 expression may aggravate the endothelial injury of diabetic nephropathy. Kidney Int, 2015. 87(2): p. 382–95.

14. Zhang, H., Z. Chen, and X. Wang, Differentiated serum levels of Kruppel-like factors 2 and 4, sP-selectin, and sE-selectin in patients with gestational diabetes mellitus. Gynecol Endocrinol, 2022. 38(12): p. 1121–1124.

15. Gao, J., et al., Endothelial Kruppel-like factor 2/4: Regulation and function in cardiovascular diseases. Cell Signal, 2025. 130: p. 111699.

16. Boon, R.A., et al., Kruppel-like factor 2 improves neovascularization capacity of aged proangiogenic cells. Eur Heart J, 2011. 32(3): p. 371–7.

17. Sangwung, P., et al., KLF2 and KLF4 control endothelial identity and vascular integrity. JCI Insight, 2017. 2(4): p. e91700.

18. Hamik, A., et al., Kruppel-like factor 4 regulates endothelial inflammation. J Biol Chem, 2007. 282(18): p. 13769–79.

19. Ng, K.W., et al., Biomarkers and point of care screening approaches for the management of preeclampsia. Commun Med (Lond), 2024. 4(1): p. 208.

20. MacDonald, T.M., et al., Clinical tools and biomarkers to predict preeclampsia. EBioMedicine, 2022. 75: p. 103780.

21. Slehria, T., et al., A Gap in Knowledge-Sudden Death and Preeclampsia. Am J Cardiol, 2023. 202: p. 199–200.

22. Atluri, N., et al., Challenges to diagnosing and managing preeclampsia in a low-resource setting: A qualitative study of obstetric provider perspectives from Ghana. PLOS Glob Public Health, 2023. 3(5): p. e0001790.

23. Akduman, A.T., et al., Effect of tamoxifen on ovarian reserve: A randomized controlled assessor-blind trial in a mouse model. J Turk Ger Gynecol Assoc, 2014. 15(4): p. 228–32.

24. Kivioja, A., et al., Increased Risk of Preeclampsia in Women With a Genetic Predisposition to Elevated Blood Pressure. Hypertension, 2022. 79(9): p. 2008–2015.

25. Sones, J.L. and R.L. Davisson, Preeclampsia, of mice and women. Physiol Genomics, 2016. 48(8): p. 565–72.

26. Dupont, V., et al., Impaired renal reserve contributes to preeclampsia via the kynurenine and soluble fms-like tyrosine kinase 1 pathway. J Clin Invest, 2022. 132(20).

27. Huang, X., et al., sFlt-1-enriched exosomes induced endothelial cell dysfunction and a preeclampsia-like phenotype in mice. Cytokine, 2023. 166: p. 156190.

28. Yang, C., et al., Long-Term Impacts of Preeclampsia on the Cardiovascular System of Mother and Offspring. Hypertension, 2023. 80(9): p. 1821–1833.

29. Wojczakowski, W., et al., Preeclampsia and Cardiovascular Risk for Offspring. J Clin Med, 2021. 10(14).

30. Turbeville, H.R. and J.M. Sasser, Preeclampsia beyond pregnancy: long-term consequences for mother and child. Am J Physiol Renal Physiol, 2020. 318(6): p. F1315–F1326.

31. Brown, M.C., et al., Cardiovascular disease risk in women with pre-eclampsia: systematic review and meta-analysis. Eur J Epidemiol, 2013. 28(1): p. 1–19.

32. Davisson, R.L., et al., Discovery of a spontaneous genetic mouse model of preeclampsia. Hypertension, 2002. 39(2 Pt 2): p. 337–42.

33. Ssengonzi, R., et al., Endothelial Nitric Oxide synthase (eNOS) in Preeclampsia: An Update. J Pregnancy Child Health, 2024. 6.

34. Kim, S., et al., Alleviation of preeclampsia-like symptoms through PlGF and eNOS regulation by hypoxia- and NF-kappaB-responsive miR-214-3p deletion. Exp Mol Med, 2024. 56(6): p. 1388–1400.

35. Zhou, Z., et al., Targeted polyelectrolyte complex micelles treat vascular complications in vivo. Proc Natl Acad Sci U S A, 2021. 118(50).

